# A widespread NADase domain links bacterial immunity with human TEP1

**DOI:** 10.64898/2026.09.04.749371

**Authors:** Wendy Le Mouëllic, Virginie Borges Cardoso, Jessie Kulsuptrakul, Julie Baltenneck, Anna Kanevskaya, Lucie Etienne, Francois Rousset

## Abstract

Recent discoveries on bacterial immunity have revealed that several protein domains involved in anti-phage defense are conserved in eukaryotes, such as SIRim, TIR, PNP and gasdermin. Bacterial immune systems therefore have the potential to illuminate fundamental biological mechanisms throughout the tree of life. Here, we report that DUF4062 domains – which we rename NIR (<u>N</u>ADase in bacterial immunity and vault ribonucleoproteins) – function in bacterial immunity against phages. We first identified NIR as an effector of Avs defense proteins, where it depletes cellular NAD^+^ to block viral infection. We then show that NIR domains are recurrently found as effectors in diverse defense systems. We use this association to uncover Vulcan and Vesta, two defense systems which trigger NAD^+^ depletion upon sensing distinct viral signals. Remarkably, NIR domains are widespread in eukaryotes where they are embedded in multiple NLR-like proteins. In particular, we identified a NIR domain with conserved NADase activity in the human TEP1 protein, a component of the telomerase complex and vault ribonucleoproteins. Together, these findings reveal an enzymatic activity shared between bacterial immunity and enigmatic eukaryotic machineries.

## INTRODUCTION

The past decade has been marked by the discovery of an outstanding number of antiphage systems in bacteria^1^. Defense systems are often made of several protein domains, such as sensors which detect phage infection, and effectors which execute cell death or dormancy to stop viral spread^2^. This architecture enables extensive domain swapping among families of defense systems: for instance, many effector domains are shared across systems, and conversely a single family of defense systems can accommodate multiple types of effectors^2,3^. This modularity can be leveraged to uncover new antiphage domains through ‘guilt-by-association’, by searching for uncharacterized protein domains in known defense systems^4–6^.

Among defense modules, effector domains can block viral propagation by targeting essential host components such as the cell membrane, nucleotides or nucleic acids^1,2^. Interestingly, NAD^+^ has emerged as a primary target of effector domains^7^. To date, five types of NAD^+^-depleting domains have been described, with TIR^8^, SIRim^9,10^ and SEFIR^11^ producing ADP-ribose (ADPR) and nicotinamide, RES domains generating ADPR-1P and nicotinamide^12^, and calcineurin-like phosphoesterase domains releasing nicotinamide mononucleotide and adenosine monophosphate^13^. NAD^+^-depleting domains are found across diverse defense systems such as Thoeris, CBASS, Argonautes and Metis^7^, further highlighting the relevance of NAD^+^ depletion for bacterial immunity.

Recent discoveries in bacterial immunity revealed that several protein domains involved in antiphage systems were previously known in the eukaryotic world^14–18^. For instance, TIR domains were initially discovered in animal Toll-like receptors (TLRs) where they mediate protein-protein interactions^19^. TIR domains were later shown to encode NADases in bacterial defenses, as well as in several immune proteins in plants^20^, in human SARM1^21^ and more recently in amoeba^22^. Conversely, bacterial defense systems are starting to reveal novel aspects of eukaryotic biology. For example, antiphage ATP nucleosidases led to the identification of a widespread family of eukaryotic effectors^6^. Likewise, the presence of SIRim domains in several antiphage systems enabled the discovery of human SIRal, a new component of the TLR immune pathway^10^. These functional and evolutionary connections highlight the potential of bacterial defense systems to drive new discoveries in eukaryotes, including in humans.

In this study, we report that NIR (previously DUF4062) is a novel NADase effector domain encoded in diverse antiphage systems. We show that this domain is found in multiple NLR-like proteins in diverse eukaryotic species, including humans. Finally, we reveal that the NADase activity of bacterial NIR domains is conserved in the human TEP1 protein, a component of the telomerase complex and vault ribonucleoproteins.

## RESULTS

### A novel NAD^+^-degrading effector domain in Avs proteins

We reasoned that novel immune effectors could be discovered by searching for variants of defense systems that encode uncharacterized protein domains in place of known effectors. We initiated our search by focusing on Avs, widespread defensive proteins gathering C-terminal repeats that sense specific viral components, a central STAND NTPase domain that mediates protein oligomerization, and a variable effector domain located at the N-terminus^23^.

While many Avs proteins encode well-characterized N-terminal effectors like nucleases or SIRim NADases, others contain an effector of unknown function^23^. Sequence analysis of an Avs protein encoding a putative CMP hydrolase from *Pseudomonas chlororaphis* (*Ps*Avs2; Refseq protein WP_273864396.1) revealed that the effector domain matches the domain of unknown function DUF4062 (Pfam PF13271) (**Fig. 1A**). To get insights into its function, we predicted the structure of the *Ps*Avs2 DUF4062 domain (*Ps*Avs2^DUF4062^; residues 1-189) with AlphaFold3^24^ and searched for structural homologues in the PDB90 database using the DALI server^25^. In line with the initial domain annotation made by Gao *et al.*^23^, a CMP-hydrolase was retrieved among the top hits. Other hits included diverse ribosyltransferases, and interestingly, many TIR domains (**Fig. 1B**). *Ps*Avs2^DUF4062^ shows high structural similarity with the TIR domain from TIR-APAZ (PDB:8qlo), an NADase belonging to the SPARTA defense system^26^ (**Fig. 1C**). Interestingly, the glutamate residue required for the catalytic NADase activity of TIR domains^21^ is conserved in *Ps*Avs2^DUF4062^ (E108) (**Fig. 1C**). Upon co-folding, AlphaFold3 confidently places NAD^+^ within the putative catalytic pocket of *Ps*Avs2^DUF4062^ where the nicotinamide-proximal ribose of NAD^+^ interacts with residue E108 (**Fig. 1D**). Altogether, these similarities with TIR domains prompted us to hypothesize that *Ps*Avs2^DUF4062^ encodes an NADase. Nevertheless, structural phylogenetic analyses revealed that DUF4062 domains cluster separately from TIR domains (**Fig. S1**), suggesting that they form a distinct family.

**Figure 1.**
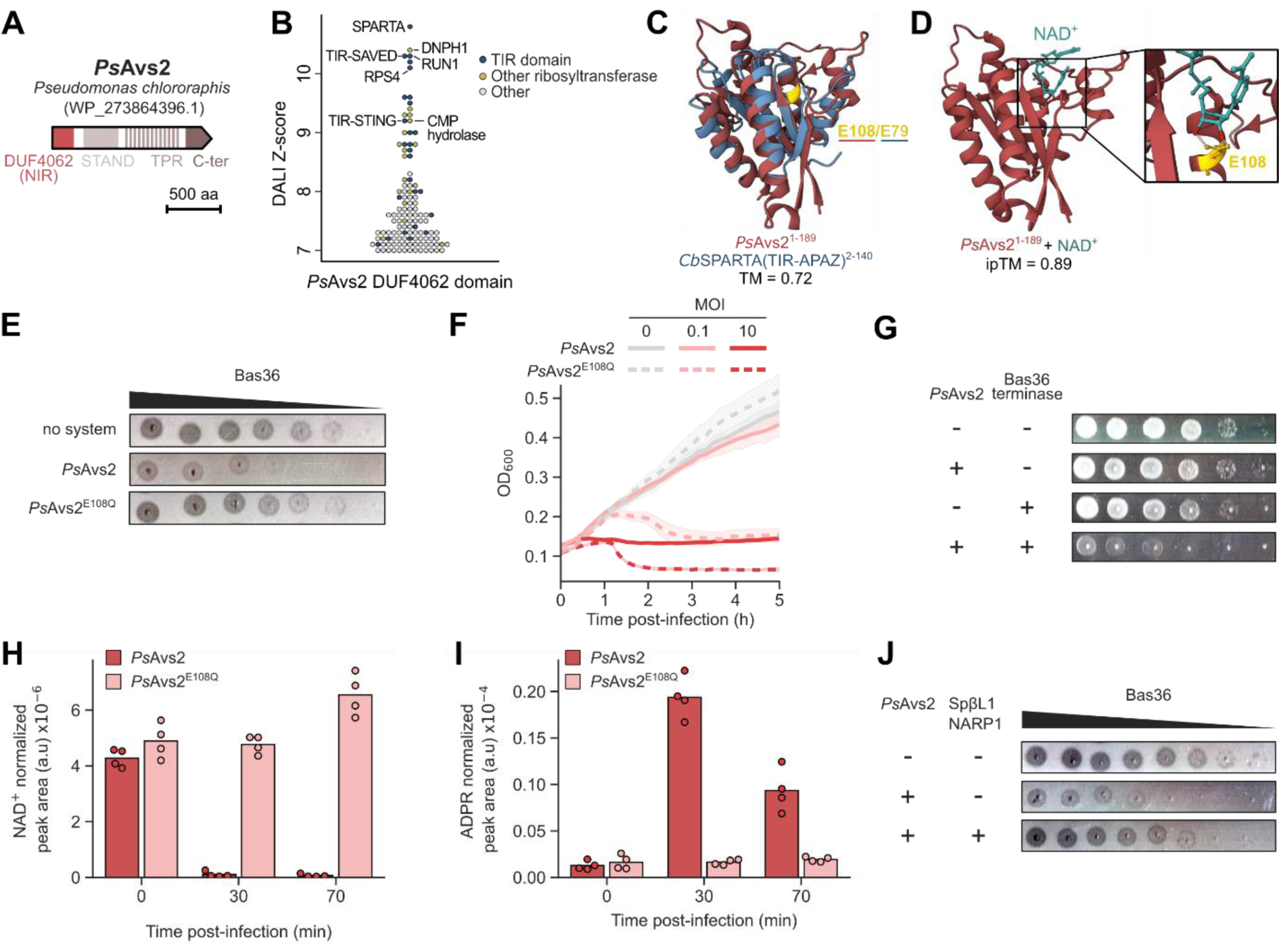
NIR (DUF4062) is a NAD^+^-degrading effector domain in Avs proteins. (**A**) Representation of a DUF4062-encoding Avs2 protein from *Pseudomonas chlororaphis*. Abbreviations: STAND, Signal Transduction ATPases with Numerous Domains; TPR, tetratricopeptide repeat. (**B**) Results of a DALI search of a predicted AlphaFold3 structure of the *Ps*Avs2 DUF4062 domain against the PDB90 database. Only hits with a Z-score > 7 are represented. (**C**) Superposition of the AlphaFold3-predicted structure of the *Ps*Avs2 DUF4062 domain with the TIR domain of the SPARTA defense system (PDB: 8qlo). Conserved glutamate residues in the active site are highlighted in yellow. (**D**) Co-folding of the *Ps*Avs2 DUF4062 domain with NAD^+^ using AlphaFold3. A zoom-in view shows the predicted interaction of E108 with the nicotinamide-proximal ribose of NAD^+^. (**E**) Plaque assay of phage Bas36 on *E. coli* strains expressing a control protein (RFP), *Ps*Avs2 or *Ps*Avs2^E108Q^. (**F**) Growth curves of *E. coli* cells expressing *Ps*Avs2 or *Ps*Avs2^E108Q^ infected with phage Bas36 at a multiplicity of infection (MOI) of 0 (uninfected), 0.1 or 10. Curves show the mean of three replicates with standard deviation displayed as a shaded area. (**G**) Ten-fold serial dilutions of *E. coli* cells expressing *Ps*Avs2 from the pLac promoter and/or the Bas36 terminase from the pTet promoter. (**H,I**) Untargeted metabolomics analysis of *E. coli* cells expressing *Ps*Avs2 or *Ps*Avs2^E108Q^ infected with phage Bas36 at a high multiplicity, showing the normalized peak area for NAD^+^ (**H**) and ADPR (ADP-ribose) (**I**). Shown is the mean of four replicates with individual datapoints overlaid. (**J**) Plaque assay of phage Bas36 on *E. coli* strains expressing *Ps*Avs2 from the pLac promoter and/or the NAD-reconstitution pathway 1 (NARP1) from phage SpβL1 from the pTet promoter.

To test its defense activity, we expressed *Ps*Avs2 in *E. coli* MG1655 ΔRM and challenged the resulting strain with the Basel collection of phages^27^. Compared to a control strain expressing a red fluorescent protein (RFP), *Ps*Avs2 provided protection against Bas34 as well as multiple phages of the *Tevenvirinae* family including Bas36 (**Fig. 1E** & **Fig. S2A**). Mutating the putative catalytic glutamate residue (E108Q) abolished defense. In liquid cultures, *Ps*Avs2 fully protected *E. coli* cells infected with a low multiplicity of infection (MOI) of Bas36, while the E108Q variant did not (**Fig. 1F**). However, *Ps*Avs2 triggered growth arrest in the presence of a high MOI of Bas36, suggesting that *Ps*Avs2^DUF4062^ targets an essential component of *E. coli* once activated. A recent study showed that an *E. coli* Avs2 protein directly senses phage terminases through its C-terminal domain^28^, which shares high structural similarity with that of *Ps*Avs2 (**Fig. S2B**). Accordingly, co-expression of the Bas36 terminase large subunit with *Ps*Avs2 was toxic to *E. coli* (**Fig. 1G**), thereby suggesting that phage terminase is an activator of *Ps*Avs2.

Because DUF4062 is structurally similar to TIR domains and requires residue E108 for defense, we tested whether *Ps*Avs2 protects against phages through NAD^+^ depletion. Analysis of polar metabolites in bacterial extracts using untargeted metabolomics revealed a marked NAD^+^ depletion during Bas36 infection in *E. coli* cells expressing *Ps*Avs2, but not in cells expressing the E108Q variant (**Fig. 1H**). This result was further confirmed by quantifying NAD^+^ concentration post-infection using a luminescence assay (**Fig. S2C**). Additionally, metabolomics analyses revealed an increase in ADP-ribose (ADPR) levels 30 minutes post-infection (**Fig. 1I**). This suggests that DUF4062 degrades NAD^+^ into ADPR and nicotinamide, similarly to TIR, SIRim and SEFIR domains in other defense systems^7^.

Phages encode NAD^+^ reconstitution pathways (NARP) to evade defense systems relying on NAD^+^ degradation such as type-I Thoeris, Dsr2 and Nehza^12,29,30^. Consistent with the NAD^+^-depleting activity of *Ps*Avs2, co-expression of the NARP1 pathway from phage SpβL1^29^ impaired *Ps*Avs2-mediated defense (**Fig. 1J**). As this pathway uses ADPR and nicotinamide to reconstitute NAD^+^, this result further supports that *Ps*Avs2^DUF4062^ catalyzes NAD^+^ hydrolysis into these products.

Altogether, our results show that DUF4062 encodes a novel NAD^+^-degrading effector domain in Avs proteins, that we rename NIR (NADase in bacterial immunity and vault ribonucleoproteins, see below).

### NIR is an immune module shared by multiple defense systems

Since bacterial defense systems are highly modular, we hypothesized that NIR domains might be encoded in additional defense systems beyond *Ps*Avs2. To gain insights into this, we used the *Ps*Avs2 NIR domain as a seed to build an HMM profile, which we searched across complete prokaryotic genomes to compute a phylogenetic tree of NIR domains (**Fig. 2**). NIR domains were widely distributed across the prokaryotic tree of life and were found in ∼8.7% of sequenced genomes. Analysis of NIR-encoding genes revealed that they are significantly enriched in the genomic vicinity of known defense genes throughout the entire tree (20.0% vs 3.9% expected by chance ; p<10^-134^, Fisher’s exact test) (**Fig. 2**, outer ring). Since defense systems tend to cluster in bacterial genomes^31,32^, this observation suggests that many, if not all, NIR-containing proteins function in prokaryotic immunity. Consistently, NIR domains were not only found in Avs proteins but also in other nucleotide-binding leucine-rich repeat (NLR)-like proteins with various C-terminal repeats. NIR domains were also found in association with a short prokaryotic argonaute and in the recently described Swarożyc^33^ and Evangelion^34^ systems. Therefore, NIR participates in diverse modular defense systems in prokaryotes.

**Figure 2.**
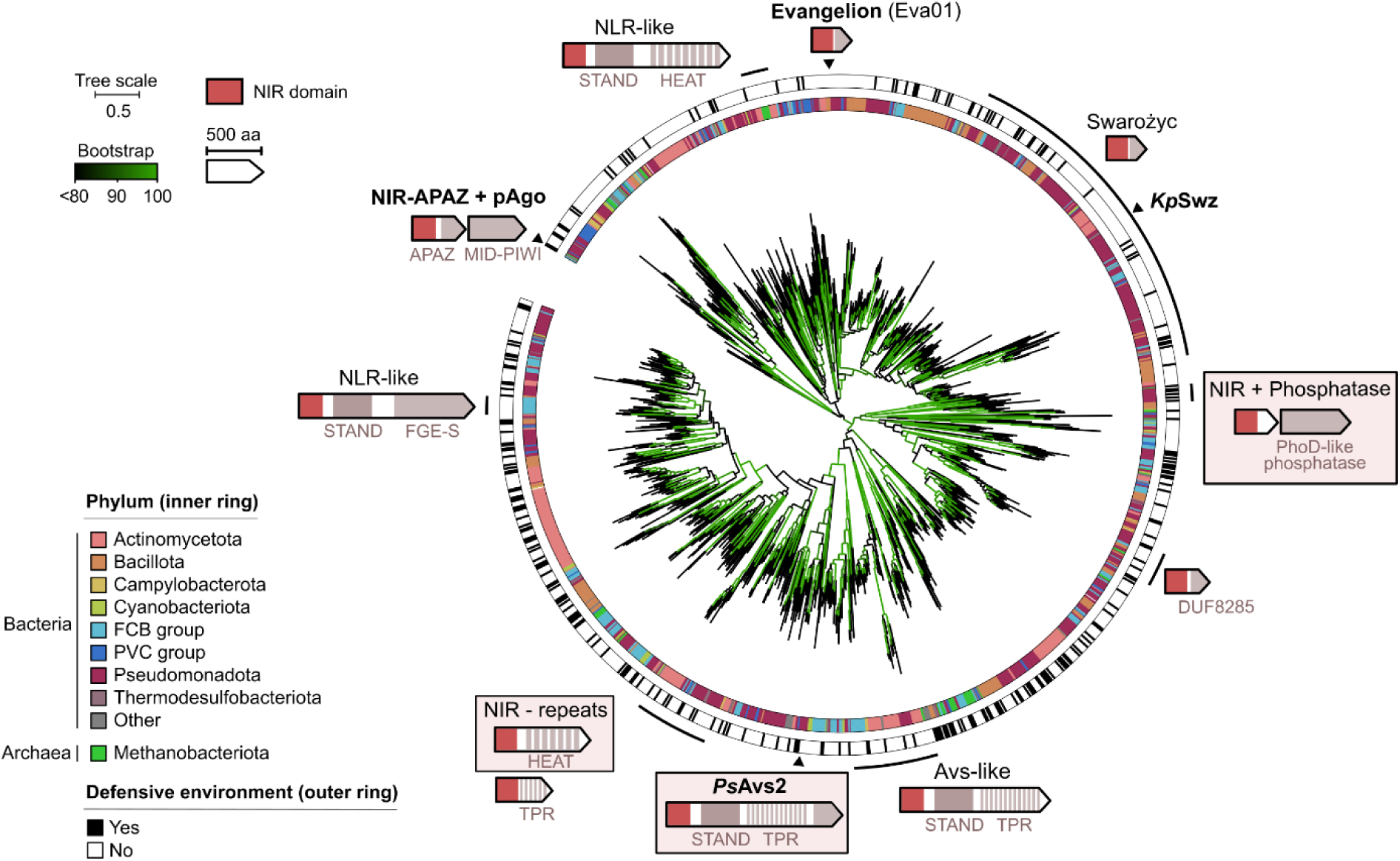
NIR is an immune module shared across defense systems. Phylogenetic tree of NIR domains across prokaryotic genomes, rooted at midpoint. Hits were clustered based on sequence identity and representative sequences were used to compute the tree. Branches are colored based on ultrafast bootstrap values^35^. Known defensive architectures are shown in bold. For each homolog shown on the tree, the presence of known defense systems within ten genes upstream or downstream was recorded (outer ring). Defense systems investigated in this study are highlighted with a box.

### Vulcan and Vesta are novel NIR-containing defense systems

We reasoned that the strong association of NIR domains with anti-phage systems could be leveraged to discover novel defense systems. We first focused on a single-protein system encoding a direct fusion between a N-terminal NIR domain and C-terminal HEAT repeats, alpha helical structures that typically function as protein-protein interaction domains^36^ (**Fig. 2** & **Fig. 3A**). A homologue from *Enterobacter roggenkampii* (RefSeq protein WP_050009967.1) provided strong defense against multiple phages of the Basel collection once expressed in *E. coli* MG1655 ΔRM, a phenotype abrogated by a single-residue mutation in the conserved catalytic glutamate residue (E98) (**Fig. 3A** and **S3A**). We named this system Vulcan, Roman god of fire. Vulcan provided robust protection at a low MOI of phage λ_vir_, but led to growth arrest at high MOI (**Fig. 3B**). Consistently, phage infection led to NAD^+^ depletion in Vulcan-expressing cells, but not in cells expressing the E98Q mutant (**Fig. 3C**). To get insights into the sensing mechanism of Vulcan, we isolated phage escapees that evade defense (**Fig. S3B**). Sequencing of seven escapees of phage λ_vir_ revealed a systematic deletion of the lambdap79 gene, which likely occurred through recombination with the lambdoid Qin prophage of *E. coli* MG1655 (**Fig. S3C**). Co-expression of lambdap79 with Vulcan led to bacterial toxicity (**Fig. 3D**), thereby revealing that lambdap79 is an activator of Vulcan. Interestingly, lambdap79 encodes a small hypothetical protein that is absent in other Basel phages blocked by Vulcan, suggesting that Vulcan can sense distinct viral proteins to trigger NAD^+^ depletion.

**Figure 3.**
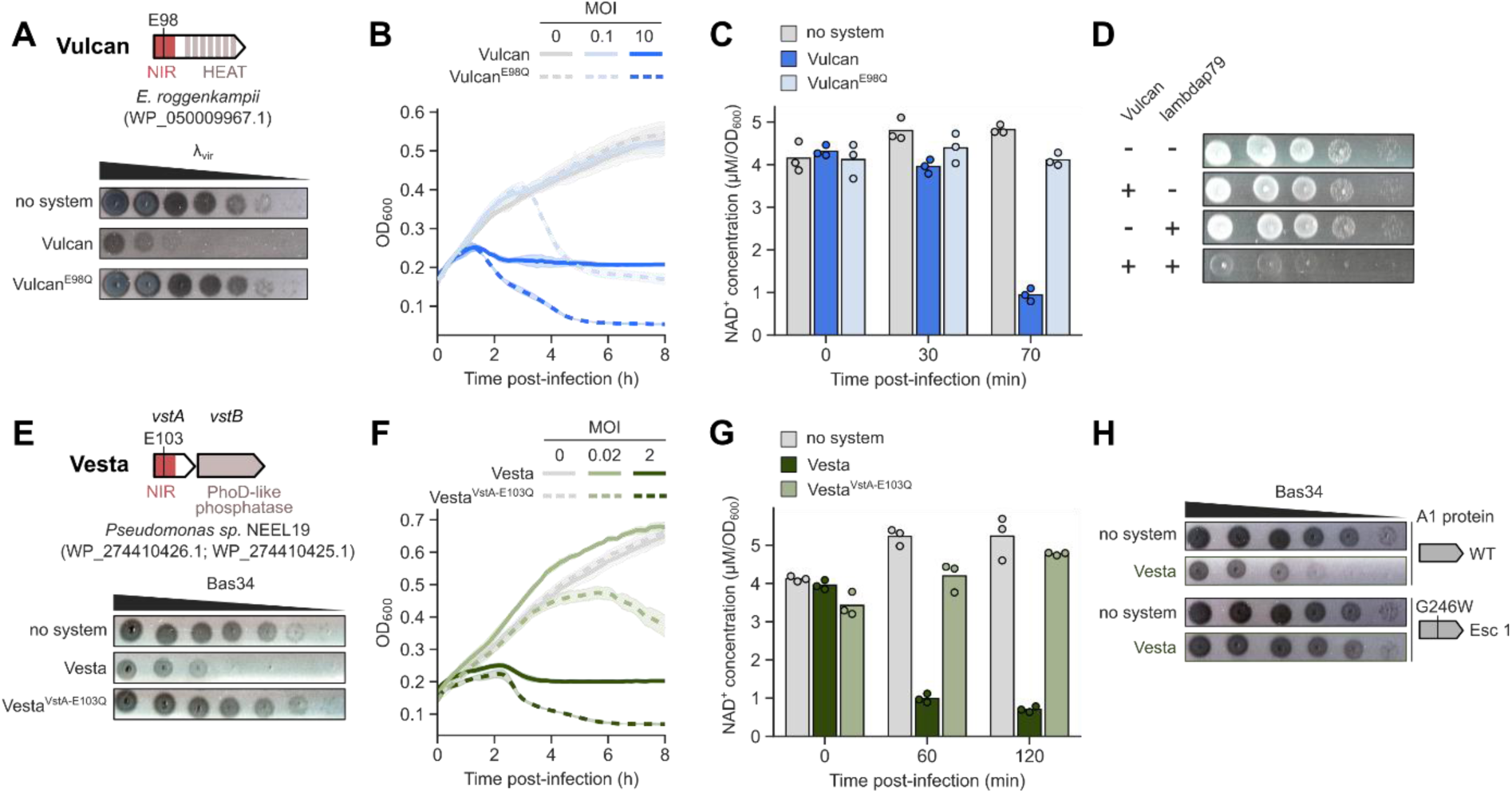
Vulcan and Vesta provide anti-phage defense through NIR-dependent NAD^+^ depletion. (**A**) Representation of a Vulcan homolog from *Enterobacter roggenkampii* and plaque assay of phage λ_vir_ on *E. coli* strains expressing a control protein (RFP), Vulcan or Vulcan^E98Q^. (**B**) Growth curves of *E. coli* cells expressing Vulcan or Vulcan^E98Q^ infected with phage λ_vir_ at a multiplicity of infection (MOI) of 0 (uninfected), 0.1 or 10. Curves show the mean of three replicates with standard deviation displayed as a shaded area. (**C**) Luminescence-based quantification of NAD^+^ concentration in *E. coli* cells expressing Vulcan or Vulcan^E98Q^ during infection with λ_vir_. (**D**) Ten-fold serial dilutions of *E. coli* cells expressing Vulcan from the pLac promoter and/or the lambdap79 gene from the pTet promoter. (**E**) Representation of a Vesta homolog from *Pseudomonas sp.* NEEL19 and plaque assay of phage Bas34 on *E. coli* strains expressing a control protein (RFP), wild-type Vesta, or Vesta with a E103Q mutation in VstA (Vesta^VstA-E103Q^). (**F**) Growth curves of *E. coli* cells expressing Vesta or Vesta^VstA-E103Q^ infected with phage Bas34 at a MOI of 0 (uninfected), 0.02 or 2. Curves show the mean of three replicates with standard deviation displayed as a shaded area. (**G**) Luminescence-based quantification of NAD^+^ concentration in *E. coli* cells expressing Vesta or Vesta^VstA-E103Q^ during infection with Bas34. (**H**) Plaque assay of wild-type Bas34 and escaper Bas34-A1^G246W^ on *E. coli* strains expressing a control protein (RFP) or Vesta.

We then focused on a two-gene system comprising a small NIR-encoding protein together with an alkaline phosphatase (**Fig. 2** & **Fig. 3E**). A homologue from *Pseudomonas sp.* NEEL19 (RefSeq proteins WP_274410426.1 and WP_274410425.1) protected *E. coli* against multiple phages, with a particularly strong defense against Bas34. This phenotype also required an intact catalytic glutamate residue (E103) in the NIR domain (**Fig. 3E** and **S3D**). We named this system Vesta, Roman goddess of hearth, home and sacred fire, and its genes *vstA* and *vstB*. Like *Ps*Avs2 and Vulcan, Vesta provided population-level protection at low – but not high – MOI (**Fig. 3F**) and led to NAD^+^ depletion during infection (**Fig. 3G**). Sequencing of Bas34-derived mutants escaping Vesta revealed non-synonymous mutations in the essential *A1* gene encoding a nuclease responsible for host genome degradation^37^ (**Fig. 3H** & **Fig. S3E**). This phenotype is reminiscent of the recently described Metis system^13^, in which the effector MisA is activated by methylated nucleotides released upon phage-induced genome degradation. Interestingly, like MisA, VstA contains a C-terminal uncharacterized domain that could function as a ligand-binding domain (**Fig. S3F**), suggesting that VstA could similarly sense a product resulting from bacterial genome degradation. Metis also encodes MisB, a HAD-family phosphatase, which is dispensable for defense but prevents toxicity in uninfected cells by degrading basal levels of methylated nucleotides. Similarly, we found that the alkaline phosphatase encoded by *vstB* is dispensable for defense (**Fig. S3D**), although *vstA* and *vstB* are strongly associated as an operon across bacterial genomes (**Fig. S3G**). Further work will be necessary to solve the full mechanism of Vesta.

Taken together, our results demonstrate that NIR is a modular effector domain found in a diversity of anti-phage systems.

### NIR domains are conserved in Eukaryotes

As many protein domains from bacterial defense systems are also found in eukaryotic proteins^14–18^, we next wondered whether NIR domains exist in eukaryotes as well. A search for NIR domains in the Eukprot v3 database^38^ (see Methods) identified thousands of hits widespread across the eukaryotic tree (**Fig. 4A** & **Table S5**). Eukaryotic NIR domains were widely distributed across phyla, with notably high prevalence in Amoebozoa (76%), Choanoflagellata (96%) and Metazoa (88%), and conversely low prevalence in Fungi (7%) and Archaeplastida (9%) (**Fig. 4B**). Notably, the vast majority of eukaryotic NIR-containing proteins consists of NLR-like proteins with the NIR domain found at the N-terminus (**Fig. 4A**, outer ring). In mammals, NIR domains are found in proteins Nwd1, Nwd2, TEP1 and TTC41 (the latter being annotated as a pseudogene in humans) (**Fig. 4A & 4C**), whose biological functions are largely unknown. Note that our search also retrieved the human protein NPHP3, whose NIR domain fell just below our significance threshold and was therefore excluded from phylogenetic analyses.

**Figure 4.**
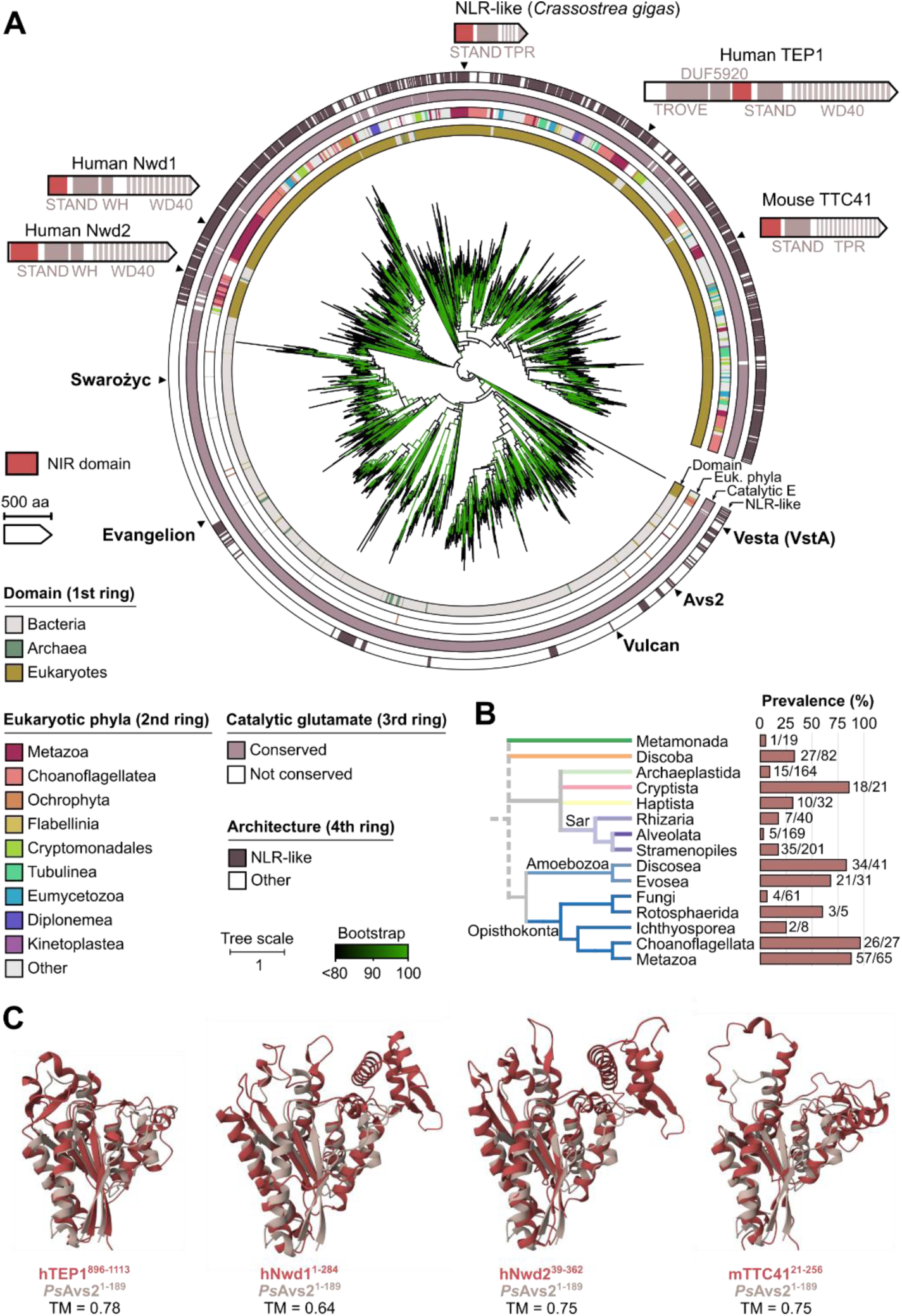
NIR domains are widespread across the tree of life. (**A**) Phylogenetic tree of NIR domains across prokaryotic and eukaryotic genomes rooted at midpoint. Hits were clustered based on sequence identity and representative sequences were used to compute the tree. Known NIR-encoding bacterial defense systems and notable eukaryotic proteins are highlighted. The ten most abundant eukaryotic phyla are annotated on the second ring. Branches are colored based on ultrafast bootstrap values^35^. Abbreviations: STAND, Signal Transduction ATPases with Numerous Domains; TPR, tetratricopeptide repeat ; TROVE, Telomerase, Ro and Vault ; WH, Winged Helix. (**B**) Prevalence of NIR domains in major eukaryotic phyla. Only phyla with at least one NIR-encoding genome are shown. Cladogram was reproduced from ref^38^. (**C**) Superposition of AlphaFold3-predicted structures of selected mammalian NIR domains with that of the *Ps*Avs2 NIR domain.

### Human TEP1 encodes an NADase

The catalytic glutamate residue essential for defense in prokaryotic NIR domains is conserved in almost all eukaryotic domains (**Fig. 4A**, third ring & **Fig. S4**), suggesting that eukaryotic homologs could have retained an enzymatic activity similar to their bacterial counterparts. To test this, we reasoned that eukaryotic NIR domains could be artificially activated upon overexpression in *E. coli* cells – in the absence of the autoinhibitory context of their native proteins – resulting in cellular toxicity, as previously shown for other effector domains^6,39^. We screened five eukaryotic domains and found that overexpression of the NIR domain from human TEP1 (hTEP1^NIR^ ; residues 896-1113) markedly reduced *E. coli* viability (**Fig. 5A**). A mutation in the putative catalytic glutamate (E1008Q in full protein) in hTEP1^NIR^ restored *E. coli* viability (**Fig. 5A**), suggesting a conserved NADase activity. In line with this hypothesis, overexpression of hTEP1^NIR^ intoxicated cells through NAD^+^ depletion, while NAD^+^ levels were maintained upon overexpression of the mutant variant (**Fig. 5B**). We therefore purified the wild-type and mutant variants of hTEP1^NIR^ and tested their NADase activity *in vitro*, by assessing their ability to process ε-NAD, an NAD^+^ analog that emits fluorescence upon cleavage. hTEP1^NIR^ showed marked ε-NAD hydrolysis activity *in vitro*, while the mutant was inactive under the same conditions (**Fig. 5C**). Ultimately, HPLC analysis of *in vitro* reactions revealed that purified hTEP1^NIR^ converts NAD^+^ into ADPR and nicotinamide (**Fig. 5D**).

**Figure 5.**
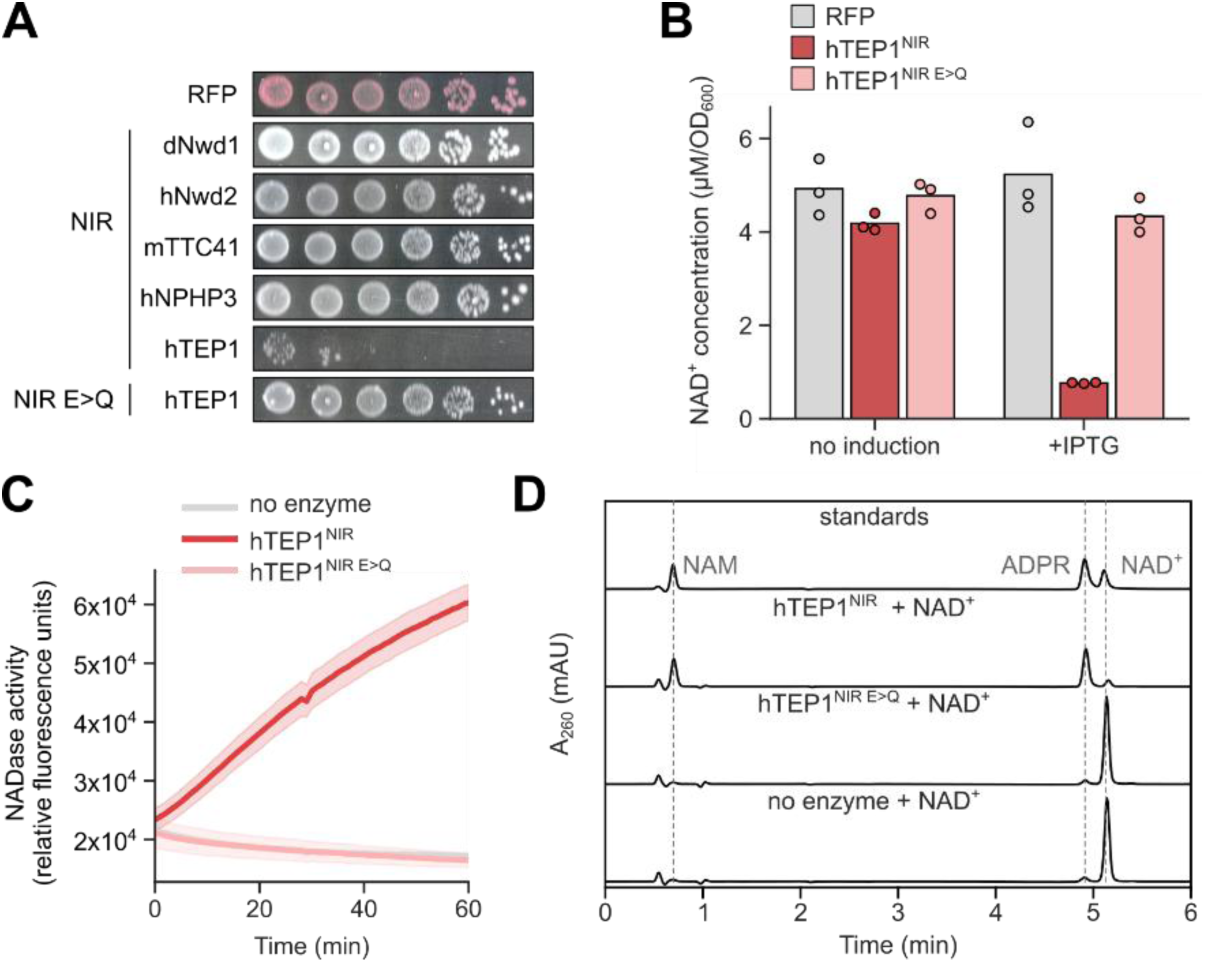
Human TEP1 contains a NIR domain with NADase activity. (**A**) Tenfold serial dilutions of *E. coli* cells expressing RFP control, wild-type or mutated (E>Q) NIR domains from selected eukaryotic proteins from a pLac promoter, spotted on LB agar supplemented with 50 µM IPTG. Species: d=*Drosophila melanogaster*; h= *Homo sapiens*; m=*Mus musculus*. (**B**) Luminescence-based quantification of NAD^+^ concentration in *E. coli* cells expressing RFP control, wild-type (hTEP1^NIR^) or mutated (hTEP1^NIR E>Q^) NIR domain from human TEP1 45 min after incubation with or without IPTG induction. (**C**) Fluorescence-based measurement of NADase activity of purified hTEP1^NIR^ and hTEP1^NIR E>Q^ domains incubated with ε-NAD. (**D**) HPLC profiles of *in vitro* reactions of purified hTEP1^NIR^ and hTEP1^NIR E>Q^ domains incubated with 500 µM NAD^+^ and profiles of chemical standards (NAD^+^: nicotinamide adenosine dinucleotide; ADPR: adenosine diphosphate ribose; NAM: nicotinamide).

Taken together, our results show that NIR domains are widespread in eukaryotic NLR-like proteins, including in several human proteins. Among them, we demonstrate that the NIR domain in human TEP1 retains the NADase activity found in bacterial anti-phage systems, thereby revealing a novel evolutionary connection between bacterial immunity and eukaryotes.

## DISCUSSION

In this study, we report NIR (DUF4062), a novel NADase domain that is shared across the tree of life. We show that the NIR domain is part of multiple antiphage systems in which it hydrolyzes NAD^+^ into ADPR and nicotinamide, leading to bacterial growth arrest and protection against phages. NIR domains therefore join the extending list of NAD^+^-degrading domains involved in bacterial immunity, which notably includes TIR^8^, SIRim^9,10^, SEFIR^11^, RES^12^ and calcineurin-like phosphoesterases^13^. The functional convergence of independent protein domains towards a NADase activity suggests that NAD^+^ depletion is a particularly efficient strategy for antiphage defense. Nevertheless, the precise effect of NAD^+^ depletion on the phage cycle remains surprisingly unclear to date.

We show that NIR domains are found in multiple NLR-like eukaryotic proteins. Among them, we highlight that the NIR domain from human TEP1 also hydrolyzes NAD^+^. Initially discovered as a component of the telomerase complex^40^, the RNA-binding protein TEP1 is also part of vault ribonucleoproteins^41^, whose biological function remains elusive more than thirty years after their discovery. While TEP1 seems dispensable for the function of the telomerase complex^42^, it is required for the association of the vault RNA with vault particles^43^. Interestingly, a recent preprint provides evidence that TEP1 is an active NADase within vault particles^33^, where it associates with poly-ADP-ribosyltransferase 4 (PARP4)^33,44^, another enzyme that uses NAD^+^ as a substrate. These indications suggest that NAD^+^ plays a central role in the yet unknown biological function of vault ribonucleoprotein particles. Future research should clarify the functional link between TEP1 and PARP4 and address whether TEP1 retains NADase activity in the telomerase complex.

Our experiments in bacteria expressing eukaryotic NIR domains, along with results obtained *in vitro* with full-length proteins^33^, show that other human NIR-containing proteins (Nwd1, Nwd2, NPHP3) do not display NADase activity under the tested conditions. These proteins might require partners to be active NADases in human cells, or might mediate other functions. This latter hypothesis is reminiscent of TIR domains, which function as NADases in bacteria, plants and amoeba but mostly mediate protein-protein interactions in animals^8^. The study of bacterial antiphage systems and plant immune proteins further revealed a role for TIR domains in the production of NAD^+^-derived signaling molecules^45–51^. A recent preprint identified a catalytic activity for animal TIR domains, including in human TLR4, which produces cyclic ADPR *in vitro*^52^. It is therefore tempting to speculate that NIR domains might have similarly undergone functional diversification over the course of evolution, with some NIR domains producing signaling molecules or functioning as protein-protein interaction modules.

## MATERIAL AND METHODS

### Bacterial strains and phages

*E. coli* K-12 MG1655 ΔRM^27^ was used for all the experiments involving bacterial defense systems and phages. *E. coli* BL21 (DE3) was used to test the activity of eukaryotic DUF4062 domains and for protein purification. *E. coli* DH5α was used as a cloning strain. Bacteria were cultivated at 37°C in LB medium supplemented with 34µg/mL chloramphenicol (Euromedex, 3886) and 50µg/mL kanamycin (Euromedex, EU0420) when required. Glucose 1% was added to the medium to cultivate strains carrying pBbA6c plasmids and *E. coli* BL21 (DE3) strains carrying pHis-TEV-derived plasmids.

Phages from the Basel collection^27^ were kindly provided by Alexander Harms’ lab and are listed in Table S1. Phages were classically amplified on *E. coli* K-12 MG1655 ΔRM at 37°C.

### Plasmids construction

*Ps*Avs2 and Vesta systems were synthesized and cloned into plasmid pBbA6c (Addgene plasmid # 35290) under the IPTG-inducible Lac promoter by Twist Bioscience. Vulcan system was synthesized as gene fragments by Twist Bioscience and assembled into plasmid pBbA6c by Gibson assembly (NEB, E2621L). Systems studied here are listed in Table S2. Eukaryotic NIR domains fused to an N-terminal His-tag with a GSG linker were synthesized as gene fragments by Twist Bioscience and cloned into a modified pHis-TEV plasmid under the T7 promoter (obtained from Christophe Rouillon). Domain sequences used in this study are listed in Table S3. Point mutations were introduced by PCR amplification using primers listed in Table S2 and S3 followed by Gibson assembly or KLD cloning (NEB, M0554S).Bas36 terminase large subunit, λ_vir_ lambdap79 and NARP1 pathway from SpβL1 phage^29^ were synthesized by Twist Bioscience and cloned into plasmid pFR66^53^ under the Tet promotor. Phage proteins used in this study are listed in Table S4.

### Plaque assays

Overnight cultures of *E. coli* K-12 MG1655 ΔRM carrying a control plasmid (pBbA6c-RFP), plasmids encoding a defense system (pBbA6c-*Ps*Avs2; pBbA6c-Vulcan; pBbA6c-Vesta) in their wild-type or mutant versions, with or without the NARP1-encoding plasmid were diluted 100-fold in 30 mL of melted LB + 0.5% agar + 5 mM CaCl_2_ + 5 µM IPTG (Vulcan) or 10 µM IPTG (other systems) (Euromedex, EU0008-B), supplemented with 0.5 µM anhydrotetracycline (aTc, Cayman Chemical, 10009542) for NARP1 experiments, poured in squared Petri dishes, dried and incubated at room temperature for 2 h. Five microliters of tenfold serial dilutions of phage suspensions were spotted on bacterial lawns. Plates were incubated for 24 h at 25°C before imaging.

### Infection in liquid medium

Overnight cultures of *E. coli* K-12 MG1655 ΔRM carrying control plasmid (pBbA6c-RFP), plasmids carrying defense systems (pBbA6c-*Ps*Avs2; pBbA6c-Vulcan; pBbA6c-Vesta) or mutated versions were diluted 100-fold in LB + 5 mM CaCl_2_ + 5 µM IPTG (Vulcan) or 10 µM IPTG (other systems) and grown at 37°C to OD_600_∼0.3 before switching to 25°C until OD_600_ reached 0.4. 180 µL of bacteria were mixed with 20 µL of LB (non-infected), 20 µL of high titer-phage suspension (high MOI) or 20 µL of 100-fold diluted phage suspension (low MOI) in a 96-well plate. Growth was followed by measuring OD_600_ every 10 min at 25°C in an Agilent BioTek Synergy H1 plate reader.

### Luminescence-based NAD^+^ quantification

For bacterial defense systems, cultures and high-MOI phage infections were performed as described in the previous section in a final volume of 1 mL in 96-deep well plates. For human TEP1 NIR domain, overnight cultures of *E. coli* BL21 (DE3) carrying plasmid control (pHis-TEV-RFP), wildtype (pHis-TEV-hTEP1^NIR^) or mutated (pHis-TEV-hTEP1^NIR^ ^E>Q^) TEP1 NIR domains, were diluted 100-fold in LB + 0.4% glucose and grown at 37°C to OD_600_∼0.5. 450 µL of cells were transferred into a 96-deep well plate and protein expression was induced by the addition of 500 µM IPTG. Cells were incubated at 37°C with agitation. At indicated times, OD_600_ was measured and 30 µL of sample were collected, mixed with 40 µL ethanol and stored at - 20°C until analysis. Samples were diluted five-fold in 100 mM phosphate buffer pH 7.55. 20 µL of sample were mixed with 20 µL of NAD/NADH-Glo^TM^ detection reagent (Promega, G9071) and incubated for 30 min at 25°C before luminescence was measured in a TECAN Infinite200 plate reader.

### Analysis of polar metabolites by LC-MS

Overnight cultures of *E. coli* K-12 MG1655 ΔRM expressing *Ps*Avs2 or *Ps*Avs2^E108Q^ were diluted 100-fold in 200 mL LB + 5 mM CaCl_2_ + 10 µM IPTG and grown at 37°C to OD_600_∼0.2 before switching to 25°C until OD_600_ reached 0.3. Cultures were infected with phage Bas36 at MOI ∼ 15 and further incubated at 25°C. Before infection, as well as 30 min and 70 min post-infection, 50 mL of culture were harvested (7 min x 4,000 g), pellets were flash-frozen and stored at -80°C until metabolites extraction. Pellets were resuspended in 500 µL of a methanol-acetonitrile-water mix (2:2:1), transferred to Lysing Matrix B tubes (MP Biomedicals, 116911500) and lysed using a FastPrep bead beater for 30 s at 6 m/s. After centrifugation (10 min x 14,000 g), supernatants were filtered through 3 kDa Amicon filters (Merck Millipore, UFC500396) (45 min x 14,000 g) and stored at -80°C until analysis.

LC-MS analyses were performed by the EMBL Metabolomics Facility as described before^54^. Briefly, LC-MS/MS analysis was performed on a Vanquish UHPLC system coupled to an Orbitrap Exploris 240 high-resolution mass spectrometer (Thermo Fisher Scientific, MA, USA) in negative and positive ESI (electrospray ionization) mode. Chromatographic separation was carried out on an Atlantis Premier BEH Z-HILIC column (Waters, MA, USA; 2.1 mm x 100 mm, 1.7 µm) at a flow rate of 0.25 mL/min. The mobile phase consisted of water:acetonitrile (9:1, v/v; mobile phase phase A) and acetonitrile:water (9:1, v/v; mobile phase B), which were modified with a total buffer concentration of 10 mM ammonium acetate (negative mode) and 10 mM ammonium formate (positive mode), respectively. The aqueous portion of each mobile phase was pH-adjusted (negative mode: pH 9.0 via addition of ammonium hydroxide; positive mode: pH 3.0 via addition of formic acid). The following gradient (20 min total run time including re-equilibration) was applied (time [min]/%B): 0/95, 2/95, 14.5/60, 16/60, 16.5/95, 20/95. Column temperature was maintained at 40°C, the autosampler was set to 4°C and sample injection volume was 5 µL. Analytes were recorded via a full scan with a mass resolving power of 120,000 over a mass range from 60 - 900 *m/z* (scan time: 100 ms, RF lens: 70%). To obtain MS/MS fragment spectra, data-dependent acquisition was carried out (resolving power: 15,000; scan time: 22 ms; stepped collision energies [%]: 30/50/70; cycle time: 900 ms). Ion source parameters were set to the following values: spray voltage: 4100 V (positive mode) / - 3500 V (negative mode), sheath gas: 30 psi, auxiliary gas: 5 psi, sweep gas: 0 psi, ion transfer tube temperature: 350°C, vaporizer temperature: 300°C. All samples were measured in a randomized manner. Pooled quality control (QC) samples were prepared by mixing equal aliquots from each processed sample. Multiple QCs were injected at the beginning of the analysis in order to equilibrate the analytical system. For determination of background signals and subsequent background subtraction, an additional processed blank sample was recorded. Data was processed using MS-DIAL 4.9^55^ and raw peak intensity data was exported for relative metabolite quantification. Level 1 feature identification was based on the in-house EMBL metabolomics library (EMBL-MCF 2.0^54^) using accurate mass, isotope pattern, MS/MS fragmentation, and retention time information with a minimum matching score of 80%.**Isolation and sequencing of escaper phages**

Putative escaper phages were retrieved from clear single plaques from plaque assays or using double-layer plaque assays as described previously^56^. Briefly, overnight cultures of *E. coli* K-12 MG1655 ΔRM carrying plasmids encoding Vulcan or Vesta were diluted 100-fold and grown in LB + 5 mM CaCl_2_ + 10 µM (Vesta) or 20 µM (Vulcan) IPTG at 37°C to OD_600_∼0.3. 100 µL of bacteria were mixed with 100 µL of phage dilutions and incubated for 10 min at room temperature. Bacteria-phage suspensions were mixed with melted LB 0.3% agar + 5 mM CaCl_2_ + 10/20 µM IPTG and poured onto a LB 1.5% agar plate. Plates were incubated for 24 hours at 25°C. Individual plaques of putative escapers were diluted in 90 µL LB and incubated for 1 hour at room temperature with frequent vortexing. Samples were centrifuged (10 min x 3,200 g) and supernatants were used to further amplify phages on *E. coli* K-12 MG1655 ΔRM. Defense of the corresponding system against initial or escaper phages was tested by plaque assay. The genome of confirmed escapers was purified as followed: 360 µL of high titer-phage suspensions were mixed with 1 µL DNAse I (NEB, M0303, 2U/µL) + 40 µL DNAse Buffer 10X (NEB, M0303) + 1 µL RNAase A (NEB, T3018, 20mg/mL) and incubated for 1 hour at 37°C. 20 µL of EDTA 0.45 M pH 8 + 10 µL of proteinase K (NEB, P8107S, 20mg/mL) were added before incubation at 56°C for 1.5 h. Samples were mixed with 800µL of gDNA binding buffer (NEB, T3014L) before loading on column and purification using Monarch® Spin PCR & DNA Cleanup Kit (NEB, T1130L) following manufacturer’s instructions. Purified genomes were sequenced by Oxford Nanopore sequencing (MicroSynth).

### Toxicity assays

To test putative phage activators, overnight cultures of *E. coli* K-12 MG1655 ΔRM coexpressing *Ps*Avs2, Vulcan or control RFP and corresponding activator (Bas36 terminase large subunit, lambda79p or control GFP) were serially diluted and spotted on LB agar + 10 µM (*Ps*Avs2) or 5 µM (Vulcan) IPTG + 0.5 µM aTc. Plates were incubated for 24 h at 25°C before imaging.

The selection of eukaryotic NIR domains tested for toxicity in *E. coli* was made based on the human proteins identified in the phylogenetic tree in Fig. 4. The human Nwd1 protein carries a truncated N-terminus that impacts the NIR domain, while TTC41 is annotated as a pseudogene in humans, with multiple stop codons dispersed throughout the coding sequence, including within the NIR domain. Instead, *Drosophila melanogaster* Nwd1 and *Mus musculus* TTC41 were respectively used. The NIR domain of human NPHP3 (Nephrocystin-3) fell just below the statistical threshold in the phylogenetic analysis presented in Fig. 4 but was included in this assay. Single colonies of *E. coli* BL21 carrying a control plasmid (pHis-TEV-RFP) or a NIR domain were diluted in 100 µL LB + 1% glucose and grown for 6 h at 37°C. Cultures were serial diluted, spotted on plates of LB agar + 1% glucose (control) or 50 µM IPTG and incubated for 24 h at 37°C before imaging.

### Protein purification

Overnight cultures of *E. coli* BL21 (DE3) carrying pHis-TEV purification plasmids were diluted 100-fold in 400 mL LB 0.4% glucose and grown at 37°C until OD_600_∼0.6. Cultures were incubated for 45 min on ice before induction of protein expression by addition of 500 µM IPTG. After overnight incubation at 20°C, cultures were centrifugated, and pellets were kept at -80°C before protein purification using NEBExpress Ni-NTA Magnetic Beads kit (NEB, S1423L), following manufacturer’s instructions. Briefly, 50 mL-culture pellets were resuspended in 1 mL binding buffer (20 mM sodium phosphate, 300 mM NaCl, 10 mM imidazole, pH 7.4), transferred to Lysing Matrix B tubes (MP Biomedicals, 116911500) and lysed using FastPrep bead beater for 30 s at 6m/s. Samples were centrifuged (10 min x 12,000 g) and supernatants were mixed to 50 µL Ni-NTA Magnetic Beads before incubation at 4°C for 30 min with rotation. Beads were washed three times with wash buffer (20 mM sodium phosphate, 300 mM NaCl, 20 mM imidazole, pH 7.4). Proteins were eluted with elution buffer (20 mM sodium phosphate, 300 mM NaCl, 500 mM imidazole, pH 7.4) and analyzed by SDS-PAGE and Coomassie staining. Buffer was exchanged to 100 mM sodium phosphate + 200 mM NaCl pH 7.55 using 10 kDa Amicon filters (Merck, UFC501096) and glycerol was added to a final concentration of 35% before storage at -20°C.

### In vitro ε-NAD hydrolysis assay

*In vitro* reactions were prepared in 100 mM phosphate buffer pH 7.55 by mixing 6 µM of purified human TEP1 NIR domain (wildtype or E1008Q) or buffer as control, and 100 µM of ε-NAD (LGC, TRC-N407735) in a final volume of 50 µL. Samples were pre-incubated for 15 min at 37°C before ε-NAD was added. Samples were transferred to black-bottom 96-well plates and fluorescence (excitation=300nm, emission=410nm) was recorded every 5 min for 1 h at 37°C on a TECAN Infinite200 plate reader.

### HPLC

In vitro reactions were prepared in 100 mM phosphate buffer pH 7.55 by mixing 6 µM of purified human TEP1 NIR domain (wildtype or E1008Q) or buffer as control, and 500 µM of NAD^+^ (ITW reagents A1124,0005) in a final volume of 30 µL. Samples were pre-incubated for 15 min at 37°C before addition of NAD^+^, and reactions were incubated for 1 h at 37°C after NAD^+^ addition. Reactions were supplemented with acetonitrile (Fisher Chemical) to a final concentration of 60%, centrifuged (5 min x 10,000 g at 4°C) and supernatants were kept for analysis. The samples were analyzed on an Agilent 1260 Infinity II HPLC system (Agilent Technologies, USA) using a Nucleoshell HILIC column (100 x 2 mm, 2.7 µm particle size; Macherey-Nagel). The column temperature was maintained at 30 °C, and the flow rate was 0.4 mL/min. Solvent A was acetonitrile (Fisher Chemical), and solvent B was 100 mM ammonium acetate (Sigma-Aldrich) in water, pH 5.0. The separation was performed using the following gradient: 0 to 1 min, 78% A and 22% B; 1 to 5 min, a linear increase of B from 22% to 60%; 5 to 7 min, 60% B. The column was then washed for 15 min and returned to the initial conditions. NAD^+^, ADPR (Merck, A0752), and nicotinamide (Merck, 72340) standards were analyzed under the same chromatographic conditions.

### Structure-based tree of DUF4062, TIR domains and other ribosyltransferases

In order to structurally compare DUF4062 with known ribosyltransferases, we first generated Alphafold3^24^ structures of DUF4062 domains from *Ps*Avs2 (WP_273864396.1, residues 1-189), human TEP1 (Uniprot Q99973, residues 896-1113) and human NPHP3 (Uniprot Q7Z494, residues 298-453) as well as TIR domains from human SARM1 (Uniprot Q6SZW1, residues 563-700) and *C. granulosa* TIR-STING (Uniprot A0A1H2ZXL9, residues 1-150). All AF3-generated structures were searched against the AFDB50^57^ database using the Foldseek^58^ server in TM-align mode. All hits with a TM-score greater than 0.5 were selected and their domain structure was extracted using hits coordinates provided by Foldseek. A structural tree based on TM scores was generated using FoldTree^59^. The tree was visualized using iTOL^60^ and leaves were annotated based on their highest-scoring Pfam using hmmscan^61^.

### Phylogenetic analyses

#### Sequence databases

Protein sequences from 41,540 complete bacterial genomes and 600 complete archaeal genomes were downloaded from Refseq in August 2024. Known defense systems were predicted using DefenseFinder (v2.0.0)^62^. Bacterial and archaeal sequences were separately filtered for redundancy using the clusthash function from MMseqs2^63^ (identity>99% and identical length). Representative non-redundant proteins were combined into a database of 49,315,157 sequences (hereafter *prokaryotic NR database*). Proteins from 993 eukaryotic genomes were downloaded from the Eukprot v3 database^38^. When several isoforms were annotated for the same protein, only the longest isoform was kept.

#### Prokaryotic search

To search for prokaryotic DUF4062, a seed HMM profile was generated as follows: homologs of the DUF4062 domain (residues 1-189) of *Ps*Avs2 (WP_273864396.1) were searched into the prokaryotic NR database using the *search* function of MMseqs2 with 2 iterations. Hits passing selection threshold (evalue<10^-3^ and query coverage >70%) were selected, their domain sequences were collected with 10 additional residues on each side and clustered with the easy-cluster function of MMseqs2 (--min-seq-id 0.8 -c 0.8). Cluster representatives were aligned with MAFFT^64^ and the resulting multiple-sequence alignment (MSA) was trimmed with clipkit^65^ (with default parameters) and used to build an HMM profile with the hmmbuild function of HMMER^61^.

The seed HMM profile was searched into the prokaryotic NR database with hmmsearch^61^. Hits passing selection threshold (evalue <10^-5^ and query coverage >70%) were selected and domain sequences were collected based on envelope residues. Domain sequences were aligned with FAMSA2^66^; in parallel, domains were clustered with the easy-cluster function of MMseqs2 (--min-seq-id 0.8 -c 0.8), and only cluster representatives were kept in the FAMSA2-generated MSA. The resulting MSA was manually trimmed on the N- and C-terminus, and a new HMM profile was generated from the trimmed MSA. This procedure was repeated twice. The final MSA was trimmed with clipkit (default parameters) and a prokaryotic phylogenetic tree was computed with IQ-TREE2^67^, with model LG+G4 selected by ModelFinder^68^. Node support was computed using 1000 iterations of the ultrafast bootstrap function^35^ in IQtree2 (option -bb 1000). Branches longer than 1.5 were discarded and the tree was recomputed with the same parameters. All trees were visualized with iTOL. Leaves were annotated as “defense-associated” when located within ten genes upstream or downstream of a known defense system.

#### Eukaryotic search

Eukaryotic DUF4062 domains are very diverse and often contain large insertions between beta sheets and alpha helices that obscure homology detection using a prokaryote-specific HMM profile. To optimize detection, we first generated a seed eukaryotic-specific HMM profile using a structure-based alignment as follows: predicted structures of DUF4062 domains from human TEP1 (Q99973, residues 892-1118), Nwd2 (Q9ULI1, residues 40-363) and TTC41P (Uniprot Q6P2S7, residues 20-252) were generated with AlphaFold3 and were searched against the AFDB50 database using Foldseek in TM-mode. Hits with a TM score greater than 0.6 were selected, and the structure of each domain was extracted from the full structure based on coordinates provided by Foldseek. Domains with a mean pLDDT score lower than 80 were discarded. Remaining domains were structurally clustered using Foldseek easy-cluster (-- tmscore-threshold 0.9). Cluster representatives were aligned using Foldmason^69^ and the resulting MSA was used to build the seed HMM profile using the hmmbuild function of HMMER^61^.

This eukaryotic seed HMM profile was searched in the Eukprot v3 database using hmmsearch. Hits passing the selection threshold (evalue < 1e-5 and query coverage > 70%) were selected, and domains were aligned with FAMSA2. In parallel, domains were clustered with the easy-cluster function of MMseqs2 (--min-seq-id 0.8 -c 0.8), and only sequence representatives were kept from the FAMSA2-generated MSA. The MSA was manually trimmed on the N- and C-terminus and a new HMM profile was generated with hmmbuild. This procedure was repeated twice, resulting in 1,751 detected eukaryotic domains clustered into 1,290 representatives.

#### Prokaryotic and eukaryotic tree

All detected prokaryotic and eukaryotic domains were combined and aligned with FAMSA2. In parallel, all domains were clustered with the easy-cluster function of MMseqs2 (--min-seq-id 0.8 -c 0.8), and only sequence representatives were kept from the FAMSA2-generated MSA. The MSA was manually trimmed on the N- and C-terminus and a final HMM profile was generated with hmmbuild. The homology search procedure was repeated once as described above using the final HMM profile against the combined prokaryotic NR database and EukProt, resulting in 3,590 sequences (1,875 prokaryotic and 1,715 eukaryotic) clustered into 2,450 representative sequences. The final MSA was trimmed with clipkit (with default parameters) and a phylogenetic tree was computed with IQtree2, with model LG+G4 selected by ModelFinder. Node support was computed using 1000 iterations of the ultrafast bootstrap function^35^ in IQtree2 (option -bb 1000).

## Acknowledgement

We thank Aude Bernheim and members of the GenoMic and LP2L teams for critical comments on the manuscript, as well as all members of the Evocure consortium for helpful discussions. We thank Bernhard Drotleff, Francois-Xavier Lehr and other members of the EMBL Metabolomics Core Facility for the acquisition and analysis of liquid chromatography-mass spectrometry data. We acknowledge support from the CBPsmn (PSMN, Pôle Scientifique de Modélisation Numérique) of the ENS de Lyon for the computing resources.

## Author contributions

W.L.M. and F.R. conceptualized the study. W.L.M., V.B.C. and J.B. performed phage-related experiments. V.B.C., J.K. and L.E. supported the development of experiments on eukaryotic domains. W.L.M. purified domains and performed biochemical assays. A.K. performed HPLC experiments. W.L.M. and F.R. performed computational analyses. F.R. supervised the study.

W.L.M. and F.R. wrote the manuscript. All authors reviewed and edited the manuscript.

