## Supplementary Figures for "A widespread NADase domain links bacterial immunity with human TEP1"

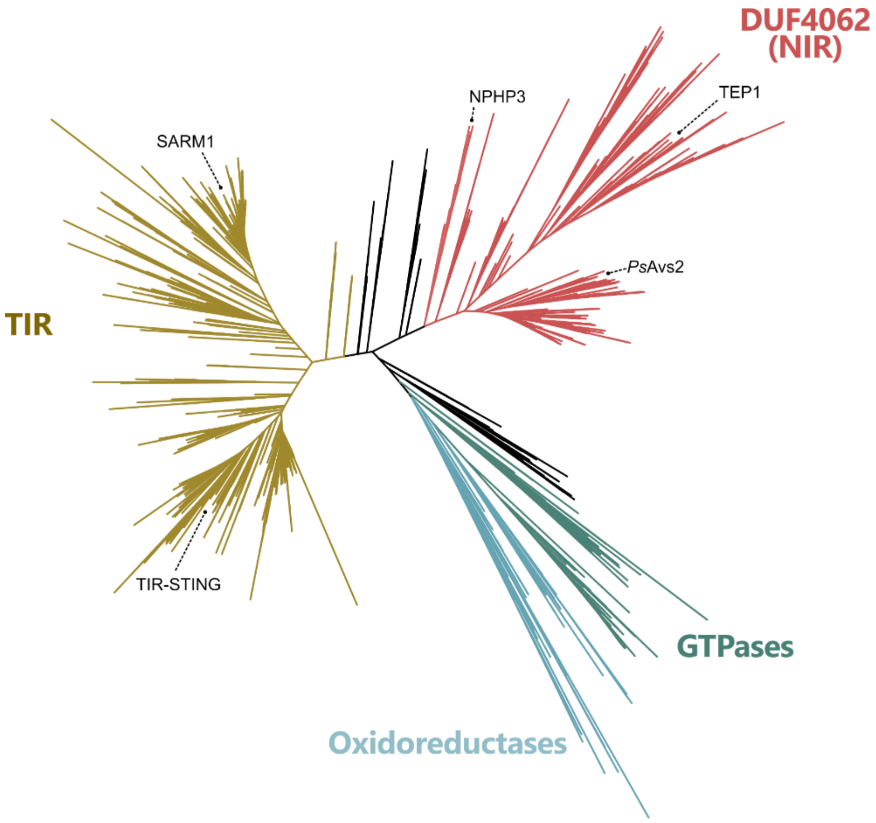


**Figure S1. Structure-based tree of DUF4062 (NIR) and TIR-like ribosyltransferases.** Predicted structures of DUF4062 domains from *Ps*Avs2 (WP_273864396.1, residues 1-189), human TEP1 (Uniprot Q99973, residues 896-1113) and human NPHP3 (Uniprot Q7Z494, residues 298-453) as well as TIR domains from human SARM1 (Uniprot Q6SZW1, residues 563-700) and *C. granulosa* TIR-STING (Uniprot A0A1H2ZXL9, residues 1-150) were generated with AlphaFold3[1] and searched against the AFDB50 database[2] in TM mode using Foldseek[3]. Hits with a TM score > 0.5 were selected and a TM-based tree was generated with FoldTree[4].

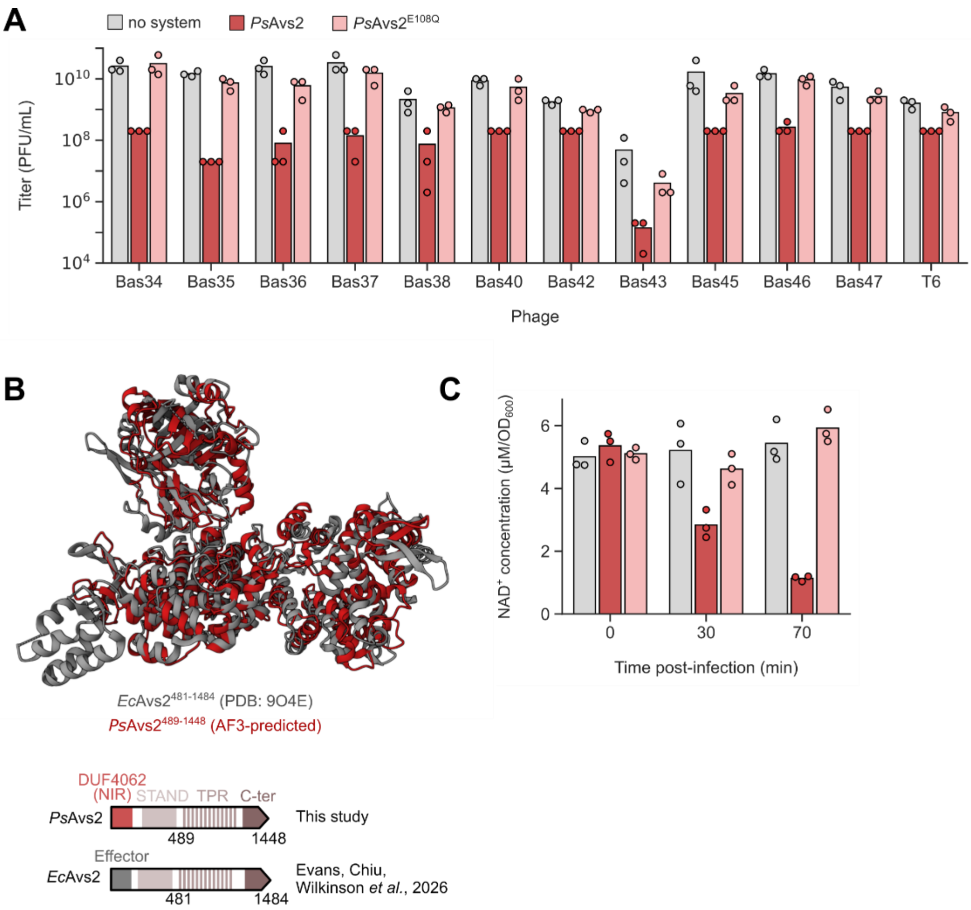


**Figure S2. The defensive activity of *Ps*Avs2.** (**A**) Plaque assays performed with phages infecting *E. coli* cells expressing RFP, *Ps*Avs2 or *Ps*Avs2$^{E108Q}$. Bars show the average of three replicates with individual datapoints overlaid. (**B**) Superposition of the AlphaFold3-predicted structure of the C-terminal region of *Ps*Avs2 with that of *Ec*Avs2 (PDB:9O4E). (**C**) Luminescence-based quantification of $NAD^+$ concentration in *E. coli* cells expressing RFP, *Ps*Avs2 or *Ps*Avs2$^{E108Q}$ during infection with phage Bas36.

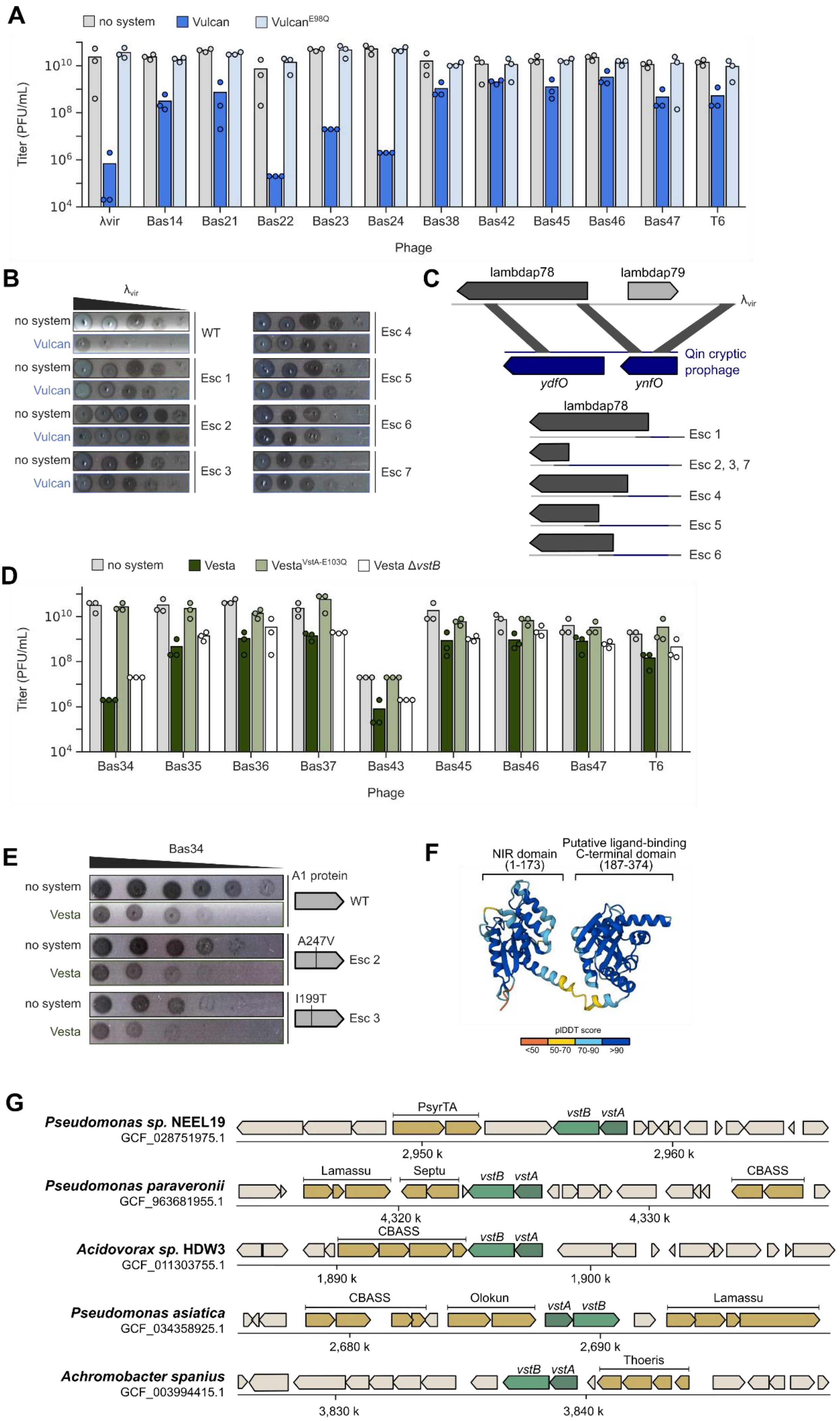

A
no system
Vulcan
VulcanE98Q
Titer (PFU/mL)
λvir
Bas14
Bas21
Bas22
Bas23
Bas24
Bas38
Bas42
Bas45
Bas46
Bas47
T6
Phage
B
λvir
no system
Vulcan
WT
Esc 1
Esc 2
Esc 3
Esc 4
Esc 5
Esc 6
Esc 7
C
lambdap78
lambdap79
λvir
Qin cryptic prophage
ydfO
ynfO
Esc 1
Esc 2, 3, 7
Esc 4
Esc 5
Esc 6
D
no system
Vesta
VestaVstA-E103Q
Vesta ΔvstB
Titer (PFU/mL)
Bas34
Bas35
Bas36
Bas37
Bas43
Bas45
Bas46
Bas47
T6
Phage
E
Bas34
no system
Vesta
A1 protein
WT
A247V
Esc 2
I199T
Esc 3
F
NIR domain (1-173)
Putative ligand-binding C-terminal domain (187-374)
pIDDT score
<50
50-70
70-90
>90
G
Pseudomonas sp. NEEL19
GCF_028751975.1
PsyrTA
vstB
vstA
2,950 k
2,960 k
Pseudomonas paraveronii
GCF_963681955.1
Lamassu
Septu
CBASS
4,320 k
4,330 k
Acidovorax sp. HDW3
GCF_011303755.1
CBASS
1,890 k
1,900 k
Pseudomonas asiatica
GCF_034358925.1
Olokun
2,680 k
2,690 k
Achromobacter spanius
GCF_003994415.1
Thoeris
3,830 k
3,840 k

**Figure S3. The defensive activity of Vulcan and Vesta.** (**A**) Plaque assays performed with phages infecting *E. coli* cells expressing RFP, Vulcan or Vulcan$^{E98Q}$. Bars show the average of three replicates with individual datapoints overlaid. (**B**) Plaque assay of wild-type and mutant phage $\lambda_{vir}$ on *E. coli* cells expressing RFP or Vulcan. (**C**) Representation of the genomic recombination of $\lambda_{vir}$ escaper mutants with the Qin prophage encoded in the genome of *E. coli* MG1655. (**D**) Plaque assays performed with phages infecting *E. coli* cells expressing RFP, Vesta or Vesta$^{VstA\text{-}E103Q}$. Bars show the average of three replicates with individual datapoints overlaid. (**E**) Plaque assay of wild-type and mutant phage Bas34 on *E. coli* cells expressing RFP or Vulcan. Non-synonymous mutations in the A1-encoding gene are represented. (**F**) AlphaFold3-predicted structure of VstA. (**G**) Genomic context of Vesta defense systems across prokaryotic genomes, showing conserved association of *vstA* and *vstB*.

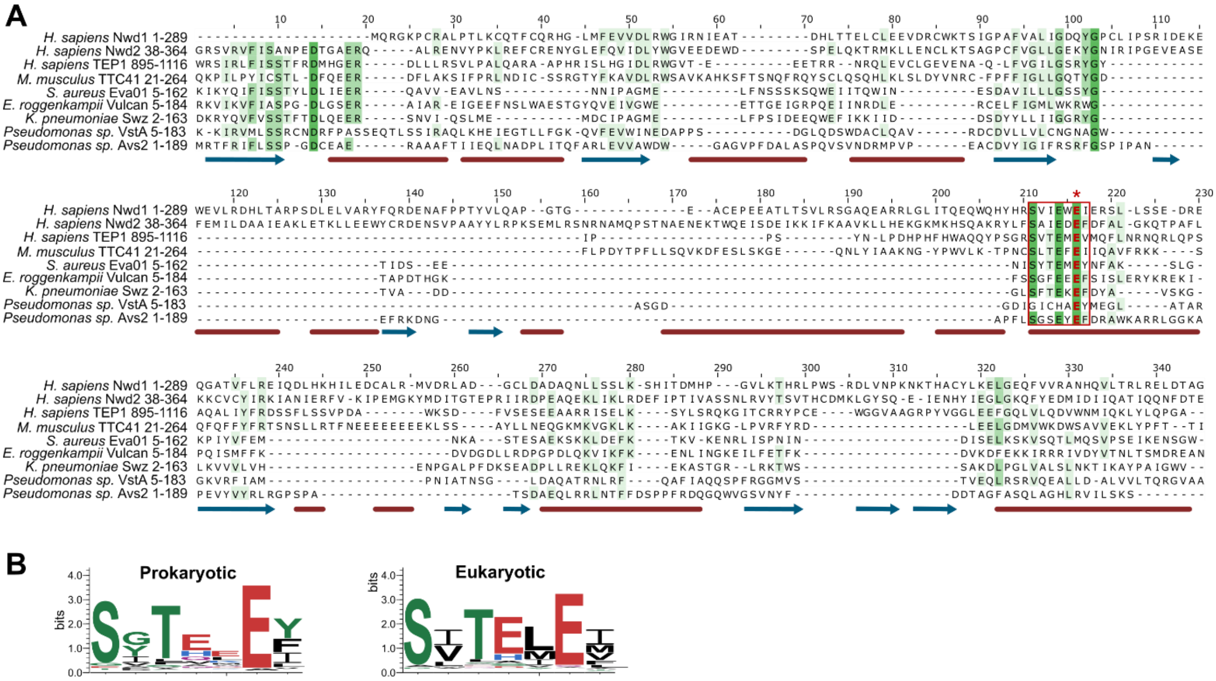

**Figure S4. Sequence analysis of eukaryotic and prokaryotic NIR domains.** (**A**) Multiple-structural alignment[5] of the NIR domains of human NWD1, human NWD2, human TEP1, mouse TTC41, and bacterial Evangelion (Eva01), Vulcan, Swarożyc, Vesta and Avs2. Predicted secondary structures are shown as blue arrow (alpha helices) and red blocks (beta barrels). The conserved SxTxxE motif shown in panel B is highlighted with a red box and the catalytic glutamate with a star. (**B**) Sequence logo of detected NIR domains in prokaryotic and eukaryotic proteins, highlighting the conserved SxTxxE motif.

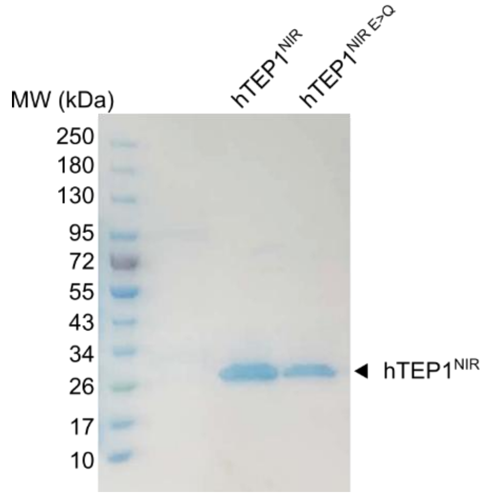


**Figure S5. SDS-PAGE analysis and Coomassie staining of wild-type and mutant hTEP1 NIR domains.** The expected molecular weight is 26.9 kDa.
